# Systematic analysis of microorganisms’ metabolism for selective targeting

**DOI:** 10.1101/2023.07.14.549016

**Authors:** Mehdi Dehghan Manshadi, Payam Setoodeh, Habil Zare

## Abstract

Selective drug targets (i.e., narrow-spectrum antibiotics) can minimize side effects of antibiotic treatments compared to broad-spectrum antibiotics due to their specific targeting of the organisms responsible for the infection. Furthermore, combating an infectious pathogen, especially a drug-resistant organism, is more efficient by attacking multiple targets. Here, we combined synthetic lethality with selective drug targeting to obtain multi-target and organism-specific potential drug candidates by systematically analyzing the genome-scale metabolic models of six different microorganisms. By considering microorganisms as targeted or conserved in one- to six-member groups, we designed 665 individual case studies. For each case, we identified single essential reactions as well as double, triple, and quadruple synthetic lethal reaction sets that are lethal for targeted microorganisms and neutral for conserved ones. As expected, the number of obtained solutions for each case depends on the genomic similarity between the studied microorganisms. Mapping the identified potential drug targets to their corresponding pathways showed the importance of key subsystems such as cell envelope biosynthesis, glycerophospholipid metabolism, membrane lipid metabolism, and the nucleotide salvage pathway. To assist validation and further investigation of our proposed potential drug targets, we introduced two sets of targets that can potentially address a substantial portion of the 665 cases. We expect that the obtained solutions provide helpful insights into designing narrow-spectrum drugs that selectively cause system-wide damage only to the target microorganisms.

## Introduction

Elimination of pathogens is the primary goal of any efficient treatment of infection. Broad-spectrum antibiotics have changed the way infectious diseases are treated. They have proven to be a precious tool in the fight against infections, and they are the most promising treatment when the exact identity of the problem-causing pathogen is unknown ^1, 2^. However, pathogens are not the only targets of broad-spectrum antibiotics. Whether administered orally or not, antibiotics can directly or indirectly influence the human microbiome, which is known as a virtual organ in the human body ^2^. The cumulative weight of the human microbiome is comparable to the liver and outnumbers human cells. The human microbiome has been the focus of many studies in recent decades for the etiology of diseases of the liver ^3, 4^, brain ^5–7^, kidney ^8, 9^, and other human organs ^10–12^. These studies suggest the critical role of the microbiome in regulating human health.

The consequences of antibiotic-induced alterations in the microbial composition include variations in microbiota functional characteristics, reduction in microbial diversity, and growth potential for the emergence of drug-resistant microorganisms ^13^. For instance, in the short term, fluoroquinolones and β-lactams decreased the diversity of microorganisms by 25% and cut the number of core phylogenetic microbiota from 29 to 12 ^14^. Additionally, broad-spectrum antibiotics, specifically tetracyclines and macrolides, might be associated with irritable bowel syndrome (IBS) development ^15^. Furthermore, the effects of antibiotics on inflammatory bowel disease (IBD) ^16, 17^, obesity-related disorders ^18^, and liver disease ^19, 20^ have been reported in the literature ^21^. On the other hand, narrow-spectrum antibiotics target the specific pathogen that causes the infection. Therefore, many studies suggest using narrow-spectrum antibiotics to overcome the problems related to taking broad-spectrum antibiotics by minimizing the alteration in a patient’s microbiome ^1, 22–25^.

Identification of selective drug targets facilitates the design of narrow-spectrum antibiotics. However, finding potential drug targets is the first step toward obtaining selective ones. Different methods for identifying potential drug targets use computational approaches ^26^. For example, genome-scale metabolic networks (GENREs), as appropriate and valuable computational tools, have significantly contributed to systems metabolic engineering ^27–31^ and *in silico* drug target identification ^32–35^. Using genome-scale metabolic models (GEMMs) as mathematical representatives of GENREs and applying constraint-based computational methods such as flux balance analysis (FBA) ^36^, numerous drug targets for *Escherichia coli*, *Helicobacter pylori*, *Mycobacterium tuberculosis, Staphylococcus aureus* ^32^, *Acinetobacter baumannii* AYE ^37^, and *Pseudomonas aeruginosa* PAO1 ^38^ have been identified. Among these studies, Lee et al. ^32^ provided selective metabolite drug targets for four pathogens using choke-point analysis ^39^. However, none of these studies has investigated the challenges of conserving one or more microorganisms while targeting the others.

According to the literature, multi-target drugs are more efficient in competing against infectious pathogens, especially drug-resistant microorganisms ^40–42^. The concept of synthetic lethality and identification of synthetic lethal sets paves the way for designing multi-target drugs. In this study, we combined the idea of synthetic lethality with selective drug targeting to obtain multi-target, microorganism-specific potential drug candidates for six different microorganisms depicted in Figure 1. As a result, for these six microorganisms we identified single essential reactions as well as double, triple, and quadruple synthetic lethal sets for rich medium with high oxygen availability, as their most resilient state. In addition to these six cases, we investigated 659 combinations between the six microorganisms generated by considering microorganisms as targeted or conserved in two-to six-member groups. Additionally, we conducted an analysis to understand the role and contribution of different pathways in the obtained drug targets. We also introduced two sets of reactions capable of handling a considerable portion of all cases. The acquired answers should be useful in helping to design narrow-spectrum antibiotics that selectively cause system-wide damage only to the target microorganisms.

**Figure 1.**
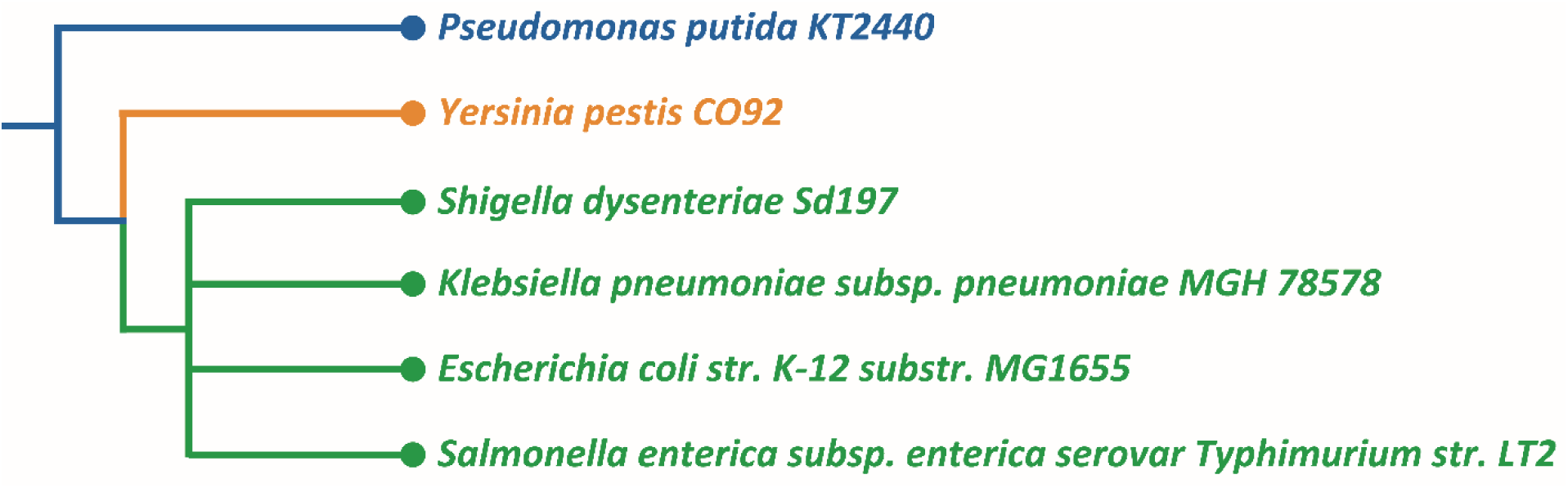
The phylogenetic tree of the six microorganisms generated by iTOL ^44^. According to this tree, *P. putida* is the most different strain among these six microorganisms. Furthermore, *S. dysenteriae*, *K. pneumoniae*, *E. coli*, and *S. enterica* stand at the same level.

The decision of choosing these six microorganisms was primarily driven by the unavailability of extensively curated models for microorganisms. Only one of these microorganisms (i.e., *Escherichia coli* str. K-12 substr. MG1655) is non-pathogenic^43^ and when comprehensive and reliable curated models for other non-pathogenic microorganisms will become available, our approach would be fundamentally applicable to include them in a similar analysis. Figure 1 depicts the phylogenetic tree of these six microorganisms generated by iTOL ^44^.

## Results

In this work, we investigated 665 cases covering all combinations of targeted and conserved microorganisms across the six models depicted in Figure 1. Among the 665 studied cases, 63 focused only on targeting different microorganisms without conserving any, while in the other 602, at least one microorganism was conserved along with other targeted microorganisms. We categorized these two groups of case studies as *with-conservation* and *without-conservation*, respectively.

Here we reported and analyzed the results of the case studies considering the constraints of the fifth step. However, both results of the fourth and fifth steps are reported in supplementary materials (See supplementary Table S1, supplementary data S1, and supplementary data S2). Our approach is based solely on *in silico* experiments, which provide confirmatory evidence for the functionality of the identified drug targets. Further experimental validation will be needed to prove the efficacy and safety of the identified solutions.

### Investigating the results of specific case studies

#### Targeting all but one microorganism

The first row of Table 1 shows the number of potential solutions for simultaneously targeting all six microorganisms. Other rows show the number of identified potential solutions for selectively targeting five out of six microorganisms while conserving the other one. Table 1 reveals that numerous possible solutions are available to kill all six microorganisms simultaneously; however, conserving even one of the microorganisms extremely reduces the number of possible solutions, which agrees with the unanimous idea that selective targeting is challenging.

**Table 1.**
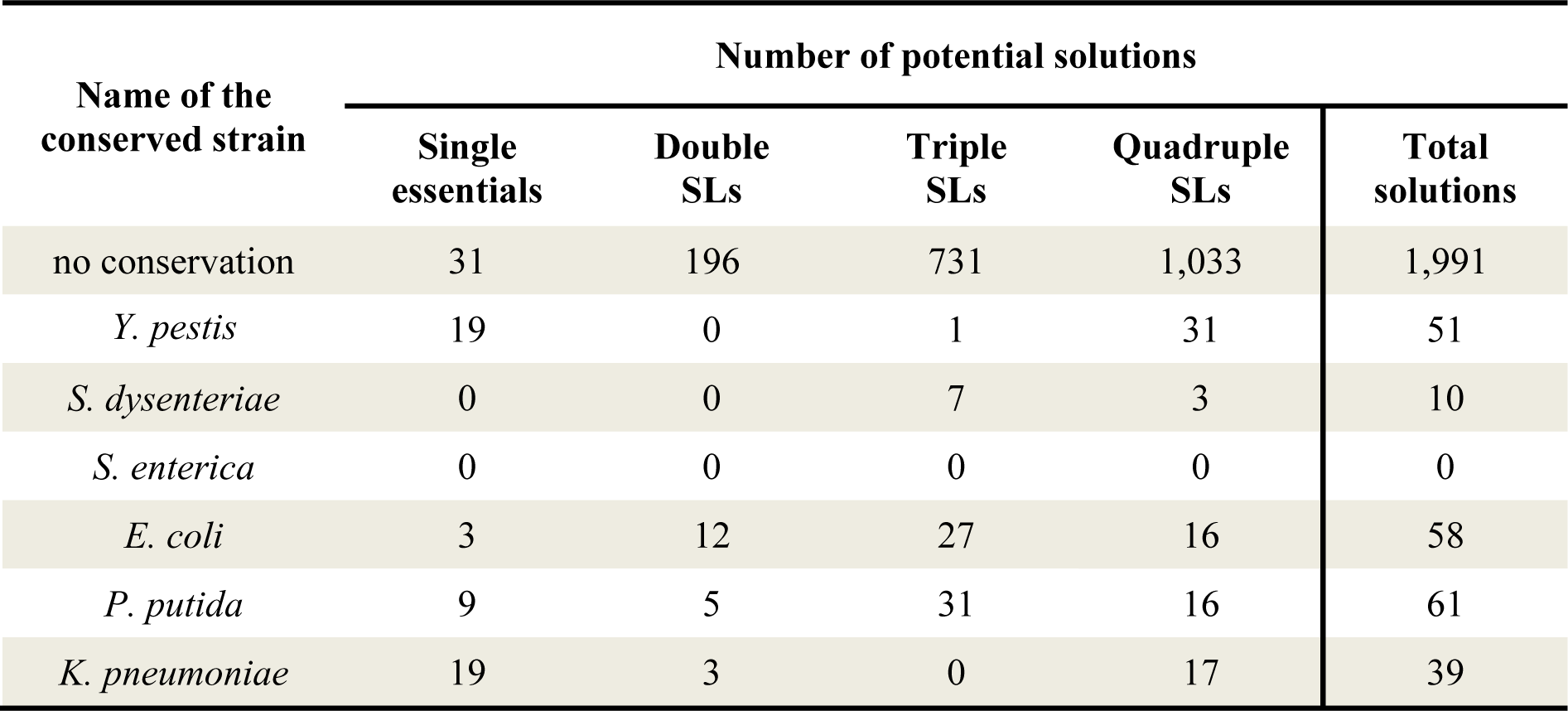
The number of potential drug targets identified for targeting five microorganisms while conserving one microorganism. Available potential solutions decreased drastically compared to the number of potential solutions for targeting all six microorganisms (the first row).

| Name of the conserved strain | Number of potential solutions |  |  |  | Total solutions |
| --- | --- | --- | --- | --- | --- |
|  | Single essentials | Double SLs | Triple SLs | Quadruple SLs |  |
| no conservation | 31 | 196 | 731 | 1,033 | 1,991 |
| *Y. pestis* | 19 | 0 | 1 | 31 | 51 |
| *S. dysenteriae* | 0 | 0 | 7 | 3 | 10 |
| *S. enterica* | 0 | 0 | 0 | 0 | 0 |
| *E. coli* | 3 | 12 | 27 | 16 | 58 |
| *P. putida* | 9 | 5 | 31 | 16 | 61 |
| *K. pneumoniae* | 19 | 3 | 0 | 17 | 39 |

According to Table 1, *S. enterica* cannot be selectively conserved. This lack of solutions is probably due to the more vulnerable state of *S. enterica* than the other microorganisms under study. We discussed this hypothesis in detail in the following sections.

#### Conserving all but one microorganism

Table 2 shows the number of the identified solutions in six cases where all but one microorganism are conserved. This table shows a relatively larger number of potential selective drug targets for *P. putida*, which suggests phylogenetic differences can lead to ease of the selective targeting of this microorganism (Figure 1).

**Table 2.** The number of potential drug targets identified for conserving five microorganisms while targeting one particular microorganism. Targeting *P. putida* has more potential solutions possibly due to its phylogenetic differences.

| Name of the targeted strain | Number of potential solutions |  |  |  | Total solutions |
| --- | --- | --- | --- | --- | --- |
|  | Single essentials | Double SLs | Triple SLs | Quadruple SLs |  |
| *Y. pestis* | 2 | 4 | 0 | 8 | 14 |
| *S. dysenteriae* | 0 | 0 | 3 | 12 | 15 |
| *S. enterica* | 2 | 0 | 0 | 0 | 2 |
| *E. coli* | 1 | 0 | 0 | 0 | 1 |
| *P. putida* | 28 | 19 | 25 | 13 | 85 |
| *K. pneumoniae* | 0 | 2 | 5 | 8 | 15 |

#### Cases with no potential solutions

There were 52 cases for which we did not find any potential solutions. In all of these cases, we were aiming for conserving microorganisms that were at the same level of the phylogenetic tree as the targeted microorganisms. All solutions for some of these cases were pretermitted after applying the strict constraints of the fifth step. However, 34 cases remained without any potential solutions, even without these constraints. In all of these cases, at least one microorganism was conserved. Only *S. enterica* was present in all 34 cases, mostly as a *conserved* strain (28 cases). Rapid-SL identified a considerably larger number of essential reactions for *S. enterica* compared to the other microorganisms under study. This issue leads to a considerable overlap between the potential solutions for *S. enterica* and other similar microorganisms (i.e., *K. pneumoniae, E. coli, and S. dysenteriae*). Consequently, gene or gene sets targeting both *K. pneumoniae* and *E. coli,* or *K. pneumoniae* and *S. dysenteriae,* were found to be deadly for *S. enterica*. The detailed results of these cases that had no solution are listed in the supplementary Table S2.

### Mapping identified solutions to pathways

In this section, we determined which pathways are more suitable for the selective targeting of microorganisms. To this aim, we marked the pathways involved based on the target reactions in each case study. We found 49 different pathways that participated in at least one potential solution. The participation rate of each pathway is reported in Supplementary Table S3. In the following, we specifically analyze some of these pathways.

#### The most targeted pathways

Cell envelope biosynthesis, glycerophospholipid metabolism, membrane lipid metabolism, and nucleotide salvage pathway are four pathways participating in all 63 cases of *without-conservation*. These pathways are related to the cell wall synthesis and the DNA replication of the microorganisms. Manipulating these pathways is the most common mechanism of available antibiotics ^45, 46^. These pathways are also the most frequently attacked in *with-conservation* cases relative to other pathways. Table 3 shows the number of *with-conservation* cases that can be accomplished by attacking these four pathways showing that no microorganism could be selectively conserved while an essential reaction from the membrane lipid metabolism pathway is targeted. In other words, these microorganisms share the same essential reactions from the membrane lipid metabolism pathway, and consequently, this pathway cannot offer any essential reaction for *with-conservation* cases.

**Table 3.** The list of pathways that appear in all 602 *with-conservation* cases. The numbers show how many cases can be accomplished by attacking the corresponding pathway. For example, 122 of 602 cases can be fulfilled by attacking the essential reactions of the Nucleotide salvage pathway.

| Pathway names | Single<br>Essentials | Double<br>SLs | Triple<br>SLs | Quadruple<br>SLs |
| --- | --- | --- | --- | --- |
| Cell envelope biosynthesis | 215 | 201 | 235 | 288 |
| Glycerophospholipid metabolism | 185 | 189 | 159 | 139 |
| Membrane lipid metabolism | 0 | 141 | 193 | 276 |
| Nucleotide salvage pathway | 122 | 181 | 220 | 190 |

#### Pathways targeted only by synthetic lethal sets

Among all solutions found, pathways such as cysteine metabolism, fatty acid biosynthesis, pyruvate metabolism, etc. were attacked only by synthetic lethal sets that include two or more reactions (i.e., none of these pathways were targeted by a single lethal reaction). In addition, no synthetic lethal set was found that maps to only one of these pathways. Supplementary Table S4 tabulates 15 pathways that have the same characteristics. Note that some synthetic lethal sets map to only one pathway, such as the pentose phosphate or glycolysis and gluconeogenesis pathway.

### Key reactions

It is valuable to find a relatively small group of reactions, the subsets of which could form different individual solutions capable of handling a considerable portion of the studied cases. Of course, obtaining the best solution for this problem is not easy and requires addressing an optimization problem that is out of the scope of this work. However, identifying near-optimal solutions is possible. We applied the greedy algorithm ^47^ to find a collection of reactions whose subsets could handle a large portion of the 665 cases. We found many four-membered combinations that can cover more than 30% of all cases. Of these, we reported a particular reaction set in Table 4 with already available drugs to target them.

**Table 4.** A group of four reactions whose subsets can fulfill 198 cases. Reaction abbreviations are the reaction IDs in the genome-scale models. The reported drug names are extracted from the DrugBank database ^58^. For each reaction (i.e., protein or gene), only one drug is reported as an example. In this table, GPR stands for the gene-protein-reaction and shows the genes associated with the reaction.

| Reactions<br>Abbreviation |  | Details |
| --- | --- | --- |
| UAGCVT | GPR | murA |
|  | Protein | UDP-N-acetylglucosamine 1-carboxyvinyltransferase |
|  | Pathway | Cell Envelope Biosynthesis |
|  | Known drug | Uridine-Diphosphate-N-Acetylglucosamine |
| 3OAS161 | GPR | fabB |
|  | Protein | 3-oxoacyl-[acyl-carrier-protein] synthase I |
|  | Pathway | Cell Envelope Biosynthesis |
|  | Known drug | Lauric acid |
| RBFSa | GPR | ribE |
|  | Protein | 6,7-dimethyl-8-ribityllumazine synthase |
|  | Pathway | Cofactor and Prosthetic Group Biosynthesis |
|  | Known drug | Dithioerythritol |
| IMPD | GPR | guaB |
|  | Protein | IMP dehydrogenase |
|  | Pathway | Purine and Pyrimidine Biosynthesis |
|  | Known drug | Inosinic Acid |

Note that the targets listed in Table 4 are combinations of single lethal reactions. As expected, the application of some of these single essential targets is limited due to drug resistance ^48–50^. Therefore, the hypothesis that attacking microorganisms with multiple targets increases the chance of overcoming drug resistance, motivated us to search for other combinations composed of only synthetic lethal sets. We obtained a combination of seven reactions whose subsets can fulfill 236 cases. Table 5 shows these seven reactions.

**Table 5.**
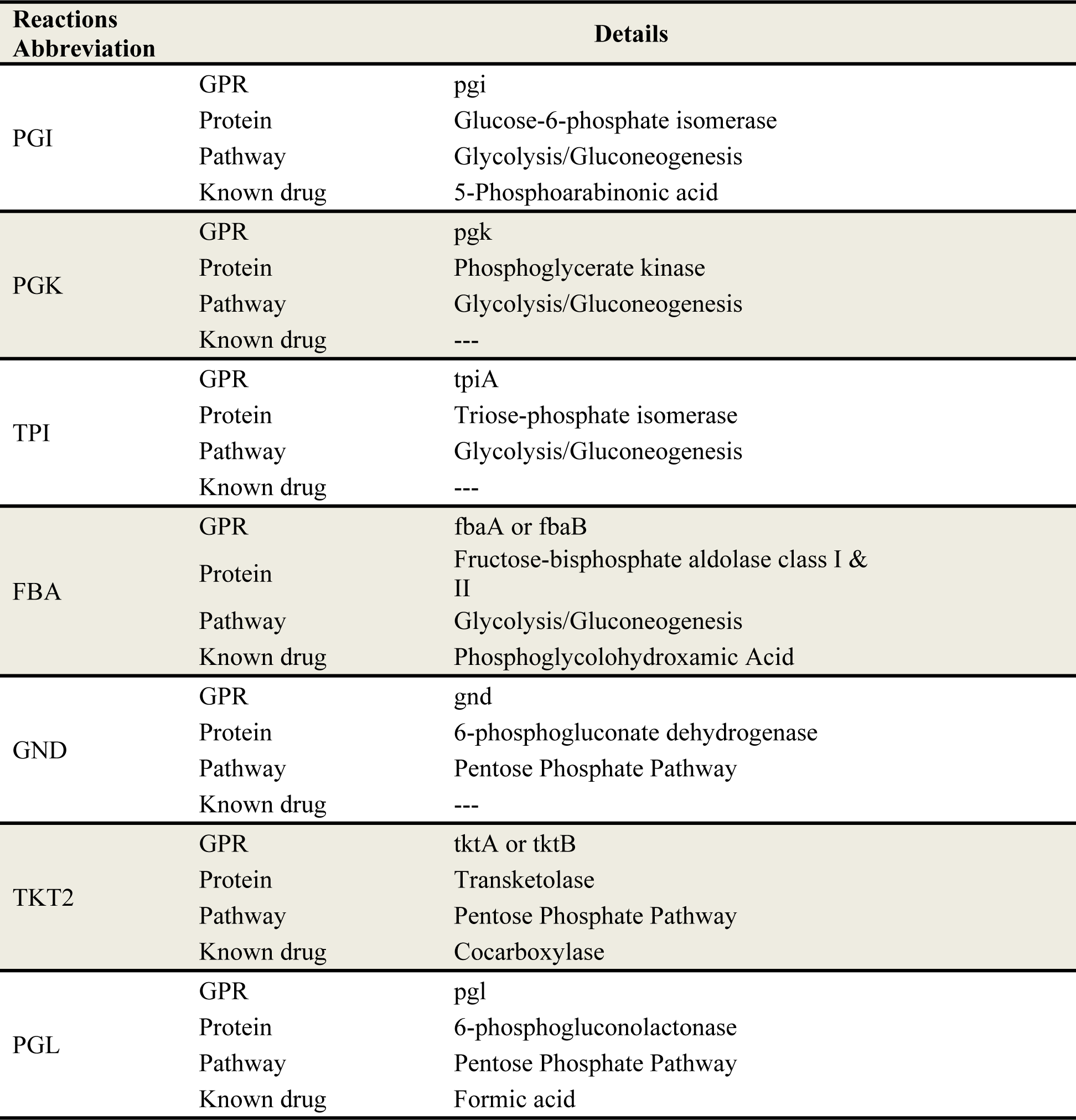
A collection of seven reactions whose subsets fulfill 236 cases. Drug names and related targets are obtained from the DrugBank database ^58^. In this table, GPR stands for the gene to protein to reaction and shows the genes associated with the reactions. These seven reactions are reported as a sample combination to address a considerable portion of all cases.

This combination is composed of three synthetic lethal sets: (1) a quadruple set of PGK, TKT2, TPI, and FBA; (2) a triple set of PGI, TPI, and GND; and (3) a double synthetic lethal set of TKT2 and PGL. These seven reactions are highly connected to the central metabolism of microorganisms.

## Discussion

Obtaining selective drug targets is exceptionally challenging compared to identifying non-selective drug targets. In this work, we investigated 665 case studies to introduce selective potential drug targets for combinations of six microorganisms. These cases include 63 *without-conservation* cases, where no microorganism is meant to be conserved, and 602 *with-conservation,* where at least one microorganism in each case is conserved.

According to the results, the cell envelope biosynthesis, glycerophospholipid metabolism, membrane lipid metabolism, and nucleotide salvage pathway contribute in all *without-conservation* cases. Also, these pathways are the targets of many potential solutions for *with-conservation* cases. These pathways are related to cell wall synthesis and DNA replication, which are reported as the main targets of many antibiotics ^45, 46^. Furthermore, our findings indicate that while the deletion of up to four reactions within certain pathways, such as cysteine metabolism, fatty acid biosynthesis, or pyruvate metabolism, may not result in a lethal set and halt the growth of the microorganism, the disruption of these pathways in conjunction with other pathways leads to cell death.

It is advantageous to identify a set of targets that can effectively address a significant number of cases. With this objective in mind, we introduced two groups of targets that have the potential to fulfill a substantial portion of the 665 cases under consideration. The first group consists of four essential reactions capable of addressing 198 cases. We also highlighted specific known drugs that target the genes associated with these reactions. As expected, the four essential reactions and their associated genes are susceptible to the development of drug resistance ^48–50^. To address this challenge, we identified a second group, including seven reactions composed exclusively of synthetic lethal sets, which can effectively target 236 cases.

One would expect that obtaining synthetic lethal sets provides more potential targets for the cases with no single lethal reactions as solutions. This hypothesis may fit the results of the fourth step, where several potential synthetic lethal solutions were obtained for 102 cases with no single lethal solution. Also, results of the fourth step show that the quadruple synthetic lethal sets generally provide much broader options to accomplish the studied cases. However, applying the strict constraints of the fifth step, which involves considering only the common reactions among the models, eliminates many triple and quadruple synthetic lethal sets. This outcome was predictable because the phylogenetic differences may facilitate the identification of selective drug targets. Consequently, limiting the analysis to common reactions in the fifth step narrows down the acceptable synthetic lethal sets and leaves 54 cases with only single targets as the solution. Thus, the results showed no considerable advantage of synthetic lethal sets over single essentials based on the solutions obtained for the fifth step.

In this work, we did not split the studied microorganisms into groups of pathogens and non-pathogens due to the lack of highly curated models of non-pathogenic microorganisms in the human gut. However, providing detailed and accurate models for these microorganisms could enable more focused investigations to identify microorganism-specific drug targets within each specific group of pathogens. By conducting targeted studies, we can gain valuable insights into the vulnerabilities unique to particular pathogens, thereby facilitating therapeutic interventions.

## Methods

We organized our study into five steps and explained each step in detail in this section.

### First step: Selecting genome-scale metabolic models

In this study, we considered a group of six microorganisms listed in Table 6. We selected these six models based on their size and their considerable number of common reactions. The selected models exhibit a remarkable level of consistency, with a total of 1048 common reactions across all six microorganisms. This shared set of reactions accounts for approximately 35% to 50% of the total reactions present in each individual model. This considerable overlap underscores the fundamental metabolic pathways and core functions that are shared among these microorganisms, despite their distinct genetic makeup and ecological niches.

**Table 6.** List of the studied microorganisms and their GEMMs.

| Microorganism Name | Model Name | No. of Genes | No. of Reactions | Ref. |
| --- | --- | --- | --- | --- |
| *Klebsiella pneumoniae* | iYL1228 | 1229 | 2262 | 59 |
| *Pseudomonas putida* | iJN1463 | 1462 | 2927 | 60 |
| *Escherichia coli* | iML1515 | 1516 | 2712 | 61 |
| *Salmonella enterica* | STM_v1_0 | 1271 | 2545 | 62 |
| *Shigella dysenteriae* | iSDY_1059 | 1059 | 2539 | 63 |
| *Yersinia pestis* | iPC815 | 815 | 1961 | 64 |

### Second step: Defining different medium conditions

Based on the availability of nutrition and oxygen, we defined four different media imposing the relevant constraints ^51^: a) a rich medium with a high oxygen uptake rate (R-H), b) a rich medium with a low oxygen uptake rate (R-L), c) a minimal medium with high oxygen uptake rate (M-H), and d) a minimal medium with low oxygen uptake rate (M-L). However, we only focused on the R-H and M-L media as the cells’ most resilient and vulnerable states, respectively.

### Third step: Identification of synthetic lethal sets

We used the concept of synthetic lethality ^52–56^ to provide potential multi-target drugs. We used Rapid-SL algorithm ^57^ to identify these synthetic lethal sets. This method deploys the depth-first-search approach to examine all potential combinations efficiently. To specify and reduce the search space for Rapid-SL, we excluded all non-gene-associated reactions, exchange reactions, demand reactions, spontaneous reactions, and diffusion reactions, which also helped us to reduce the number of not-applicable and trivial solutions. We should first obtain single essentials and synthetic lethal sets for each model. After that, we can analyze these solutions to find selective drug targets in the fourth step.

An effective potential drug target was defined as a single or synthetic lethal reaction set if it was deleterious for all four distinct medium conditions. However, there was no need to perform a lethality analysis for each medium condition separately to find common solutions. Because the solution space of the R-H medium is a subset of other medium conditions, any lethal set identified for the R-H medium is deleterious in the other medium conditions. Therefore, we performed the lethality analysis of each model just for the R-H medium condition.

### Fourth step: Identification of potential microorganism-specific and multi-target drugs

Here, we investigated various cases as groups of two to six microorganisms generated by categorizing microorganisms as targeted, conserved. Accordingly, 659 individual combinations were made, and in 57 cases, we examined the targeting of two to six microorganisms while none was conserved. In the other 602 cases, we considered at least one targeted and one conserved microorganism. Figure 2 shows a schematic of the different studied cases.

**Figure 2.**
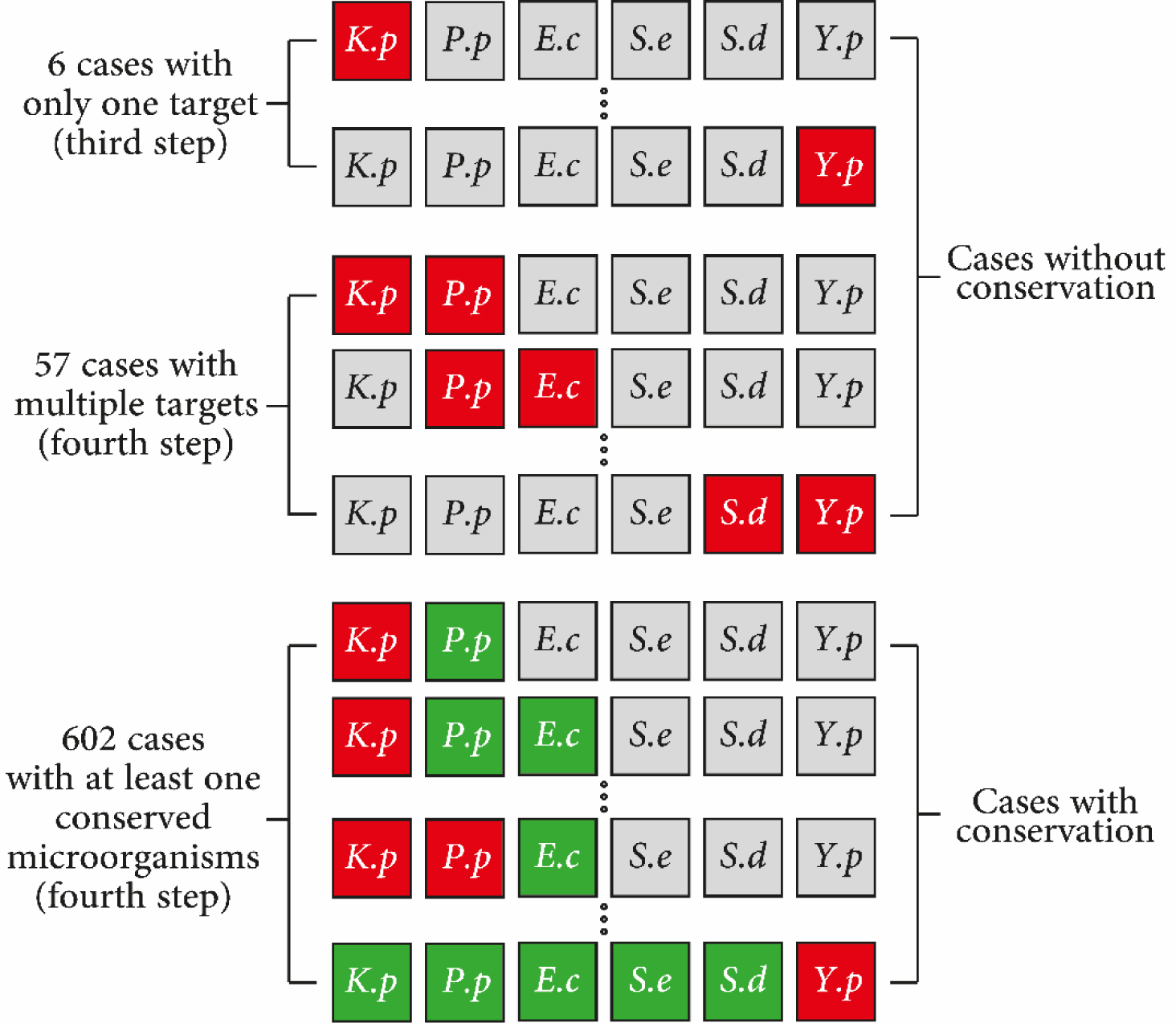
The schematic of the different studied cases. *K.p, P.p, E.c, S.e, S.d*, and *Y.p* stand for *Klebsiella pneumoniae, Pseudomonas putida, Escherichia coli, Salmonella enterica, Shigella dysenteriae,* and *Yersinia pestis*, respectively. Red, green, and gray squares represent targeted, conserved, and not included microorganisms in each case.

In this step, we checked the selectivity of each solution obtained in the third step for different case studies. We defined selective solutions as single lethal or synthetic lethal reaction sets that are deleterious for all targeted microorganisms but not for the conserved ones. However, to ensure that these selective solutions are non-lethal for all medium conditions, we tested their knockout effects for M-L medium condition of all conserved models. This constraint guarantees that conserved microorganisms survive in their most vulnerable state, and the obtained potential solution remains selective.

### Fifth step: Applying other strict constraints

There are two possibilities when a selective solution is found: (a) all target reactions are present in all conserved models, or (b) at least one target reaction is not present in one of the conserved models. In both scenarios, the knockout of target reactions is not deadly for the conserved models. However, these models only cover a fraction of their whole genome. Therefore, in the second scenario, it is probable that the target reactions and their corresponding genes are not considered in the model while they are present in the conserved microorganism’s genome. Based on this fact, we cautiously pretermitted the selective solutions obtained from the second scenario due to the lack of accurate information.

The STORMS checklist for this paper is available at: https://github.com/CSBLaboratory/SelectiveDrugTargets/blob/main/STORMS_Excel_1.03.xlsx

## Data availability

All synthetic lethality analyses, constrained models, and scripts to reproduce the results are publicly available at https://github.com/CSBLaboratory/SelectiveDrugTargets.

## Author contributions

M.D.M, P.S, and H.Z have contributed to the concept of the article. M.D.M implemented the method and obtained the results. P.S and H.Z analyzed the results. All authors wrote and reviewed the manuscript.

## References

1 Hill, C. Microbiome and infection: a case for “selective depletion”. Annals of Nutrition and Metabolism 77, 4–9 (2021).

2 Luo, Y. & Zhou, T. Connecting the dots: Targeting the microbiome in drug toxicity. Medicinal Research Reviews 42, 83–111 (2022).

3 Woodhouse, C., Patel, V., Singanayagam, A. & Shawcross, D. the gut microbiome as a therapeutic target in the pathogenesis and treatment of chronic liver disease. Alimentary pharmacology & therapeutics 47, 192–202 (2018).

4 Sarin, S. K., Pande, A. & Schnabl, B. Microbiome as a therapeutic target in alcohol-related liver disease. Journal of hepatology 70, 260–272 (2019).

5 Martin, C. R., Osadchiy, V., Kalani, A. & Mayer, E. A. The brain-gut-microbiome axis. Cellular and molecular gastroenterology and hepatology 6, 133–148 (2018).

6 Dinan, T. G. & Cryan, J. F. The microbiome-gut-brain axis in health and disease. Gastroenterology Clinics 46, 77–89 (2017).

7 Ghaisas, S., Maher, J. & Kanthasamy, A. Gut microbiome in health and disease: Linking the microbiome–gut–brain axis and environmental factors in the pathogenesis of systemic and neurodegenerative diseases. Pharmacology & therapeutics 158, 52–62 (2016).

8 Chen, Y.-Y. et al. Microbiome–metabolome reveals the contribution of gut–kidney axis on kidney disease. Journal of translational medicine 17, 1–11 (2019).

9 Shankaranarayanan, D. & Raj, D. Gut Microbiome and Kidney Disease. Clinical Journal of the American Society of Nephrology (2022).

10 Ahmadmehrabi, S. & Tang, W. W. Gut microbiome and its role in cardiovascular diseases. Current opinion in cardiology 32, 761 (2017).

11 Okuyama, Y. et al. The influence of gut microbiome on progression of overactive bladder symptoms: A community-based 3-year longitudinal study in Aomori, Japan. International Urology and Nephrology 54, 9–16 (2022).

12 Okamoto, T. et al. Altered gut microbiome associated with overactive bladder and daily urinary urgency. World Journal of Urology 39, 847–853 (2021).

13 Patangia, D. V., Anthony Ryan, C., Dempsey, E., Paul Ross, R. & Stanton, C. Impact of antibiotics on the human microbiome and consequences for host health. MicrobiologyOpen 11, e1260 (2022).

14 Panda, S. et al. Short-term effect of antibiotics on human gut microbiota. PloS one 9, e95476 (2014).

15 Villarreal, A. A., Aberger, F. J., Benrud, R. & Gundrum, J. D. Use of broad-spectrum antibiotics and the development of irritable bowel syndrome. Wmj 111, 17–20 (2012).

16 Shimodaira, Y., Watanabe, K. & Iijima, K. The risk of antibiotics and enterocolitis for the development of inflammatory bowel disease: a Japanese administrative database analysis. Scientific Reports 12, 1–8 (2022).

17 Ungaro, R. et al. Antibiotics associated with increased risk of new-onset Crohn’s disease but not ulcerative colitis: a meta-analysis. Official journal of the American College of Gastroenterology*| ACG* 109, 1728–1738 (2014).

18 Vallianou, N., Dalamaga, M., Stratigou, T., Karampela, I. & Tsigalou, C. Do antibiotics cause obesity through long-term alterations in the gut microbiome? A review of current evidence. Current Obesity Reports 10, 244–262 (2021).

19 Ferrajolo, C. et al. Antibiotic-induced liver injury in paediatric outpatients: a case-control study in primary care databases. Drug safety 40, 305–315 (2017).

20 Stine, J. G. & Lewis, J. H. Hepatotoxicity of antibiotics: a review and update for the clinician. Clinics in Liver Disease 17, 609–642 (2013).

21 Ianiro, G., Tilg, H. & Gasbarrini, A. Antibiotics as deep modulators of gut microbiota: between good and evil. Gut 65, 1906–1915 (2016).

22 Melander, R. J., Zurawski, D. V. & Melander, C. Narrow-spectrum antibacterial agents. Medchemcomm 9, 12–21 (2018).

23 Ostorhazi, E. et al. Advantage of a narrow spectrum host defense (antimicrobial) peptide over a broad spectrum analog in preclinical drug development. Frontiers in Chemistry 6, 359 (2018).

24 Alm, R. A. & Lahiri, S. D. Narrow-spectrum antibacterial agents—benefits and challenges. Antibiotics 9, 418 (2020).

25 Mondhe, M., Chessher, A., Goh, S., Good, L. & Stach, J. E. Species-selective killing of bacteria by antimicrobial peptide-PNAs. PloS one 9, e89082 (2014).

26 Chandra, N. Computational approaches for drug target identification in pathogenic diseases. Expert Opinion on Drug Discovery 6, 975–979 (2011).

27 Garcia-Albornoz, M. A. & Nielsen, J. Application of genome-scale metabolic models in metabolic engineering. Industrial Biotechnology 9, 203–214 (2013).

28 Kim, W. J., Kim, H. U. & Lee, S. Y. Current state and applications of microbial genome-scale metabolic models. Current Opinion in Systems Biology 2, 10–18 (2017).

29 Gu, C., Kim, G. B., Kim, W. J., Kim, H. U. & Lee, S. Y. Current status and applications of genome-scale metabolic models. Genome biology 20, 1–18 (2019).

30 Purdy, H. M. & Reed, J. L. Evaluating the capabilities of microbial chemical production using genome-scale metabolic models. Current Opinion in Systems Biology 2, 91–97 (2017).

31 Choi, K. R. et al. Systems metabolic engineering strategies: integrating systems and synthetic biology with metabolic engineering. Trends in biotechnology 37, 817–837 (2019).

32 Kim, T. Y., Kim, H. U. & Lee, S. Y. Metabolite-centric approaches for the discovery of antibacterials using genome-scale metabolic networks. Metabolic Engineering 12, 105–111 (2010).

33 Jerby, L. & Ruppin, E. Predicting drug targets and biomarkers of cancer via genome-scale metabolic modeling. Clinical Cancer Research 18, 5572–5584 (2012).

34 Cesur, M. F., Siraj, B., Uddin, R., Durmuş, S. & Çakır, T. Network-based metabolism-centered screening of potential drug targets in Klebsiella pneumoniae at genome scale. Frontiers in Cellular and Infection Microbiology, 447 (2020).

35 Mohite, O. S., Weber, T., Kim, H. U. & Lee, S. Y. Genome-Scale Metabolic Reconstruction of Actinomycetes for Antibiotics Production. Biotechnology Journal 14, 1800377 (2019).

36 Orth, J. D., Thiele, I. & Palsson, B. Ø. What is flux balance analysis? Nature biotechnology 28, 245–248 (2010).

37 Kim, H. U., Kim, T. Y. & Lee, S. Y. Genome-scale metabolic network analysis and drug targeting of multi-drug resistant pathogen Acinetobacter baumannii AYE. Molecular BioSystems 6, 339–348 (2010).

38 Perumal, D., Samal, A., Sakharkar, K. R. & Sakharkar, M. K. Targeting multiple targets in Pseudomonas aeruginosa PAO1 using flux balance analysis of a reconstructed genome-scale metabolic network. Journal of drug targeting 19, 1–13 (2011).

39 Yeh, I., Hanekamp, T., Tsoka, S., Karp, P. D. & Altman, R. B. Computational analysis of Plasmodium falciparum metabolism: organizing genomic information to facilitate drug discovery. Genome research 14, 917–924 (2004).

40 Korcsmáros, T., Szalay, M. S., Böde, C., Kovács, I. A. & Csermely, P. How to design multi-target drugs: target search options in cellular networks. Expert opinion on drug discovery 2, 799–808 (2007).

41 Zimmermann, G. R., Lehar, J. & Keith, C. T. Multi-target therapeutics: when the whole is greater than the sum of the parts. Drug discovery today 12, 34–42 (2007).

42 Talevi, A. Multi-target pharmacology: possibilities and limitations of the “skeleton key approach” from a medicinal chemist perspective. Frontiers in pharmacology, 205 (2015).

43 Epa, U. Escherichia coli K-12 final risk assessment: attachment I–final risk assessment of Escherichia coli K-12 derivatives. (1997).

44 Letunic, I. & Bork, P. Interactive Tree Of Life (iTOL) v4: recent updates and new developments. Nucleic acids research 47, W256–W259 (2019).

45 Kohanski, M. A., Dwyer, D. J. & Collins, J. J. How antibiotics kill bacteria: from targets to networks. Nature Reviews Microbiology 8, 423–435 (2010).

46 Murima, P., McKinney, J. D. & Pethe, K. Targeting bacterial central metabolism for drug development. Chemistry & biology 21, 1423–1432 (2014).

47 Cormen, T. H., Leiserson, C. E., Rivest, R. L. & Stein, C. Greedy algorithms. Introduction to algorithms 1, 329–355 (2001).

48 Falagas, M. E., Athanasaki, F., Voulgaris, G. L., Triarides, N. A. & Vardakas, K. Z. Resistance to fosfomycin: mechanisms, frequency and clinical consequences. International journal of antimicrobial agents 53, 22–28 (2019).

49 Zhang, Y. et al. Unraveling mechanisms and epidemic characteristics of nitrofurantoin resistance in uropathogenic Enterococcus faecium clinical isolates. Infection and Drug Resistance, 1601–1611 (2021).

50 Christaki, E., Marcou, M. & Tofarides, A. Antimicrobial resistance in bacteria: mechanisms, evolution, and persistence. Journal of molecular evolution 88, 26–40 (2020).

51 Sigurdsson, G., Fleming, R. M., Heinken, A. & Thiele, I. A systems biology approach to drug targets in Pseudomonas aeruginosa biofilm. PLoS One 7, e34337 (2012).

52 Cottarel, G. & Wierzbowski, J. Combination drugs, an emerging option for antibacterial therapy. Trends in biotechnology 25, 547–555 (2007).

53 Kaelin, W. G. The concept of synthetic lethality in the context of anticancer therapy. Nature reviews cancer 5, 689–698 (2005).

54 Klobucar, K. & Brown, E. D. Use of genetic and chemical synthetic lethality as probes of complexity in bacterial cell systems. FEMS Microbiology Reviews 42, fux054 (2018).

55 Tyers, M. & Wright, G. D. Drug combinations: a strategy to extend the life of antibiotics in the 21st century. Nature Reviews Microbiology 17, 141–155 (2019).

56 Zhang, Y. Using Synthetic-Lethal Interactions to Discover Antibacterial Drug Targets. (2022).

57 Dehghan Manshadi, M., Setoodeh, P. & Zare, H. Rapid-SL identifies synthetic lethal sets with an arbitrary cardinality. Scientific reports 12, 1–9 (2022).

58 Wishart, D. S. et al. DrugBank 5.0: a major update to the DrugBank database for 2018. Nucleic acids research 46, D1074–D1082 (2018).

59 Liao, Y.-C. et al. An experimentally validated genome-scale metabolic reconstruction of Klebsiella pneumoniae MGH 78578, i YL1228. Journal of bacteriology 193, 1710–1717 (2011).

60 Lewis, L. A., Perisin, M. A. & Tobias, A. V. Metabolic Modeling of Pseudomonas putida to Understand and Improve the Breakdown of Plastic Waste. (CCDC Army Research Laboratory Adelphi United States, 2020).

61 Monk, J. M. et al. iML1515, a knowledgebase that computes Escherichia coli traits. Nature biotechnology 35, 904–908 (2017).

62 Thiele, I. et al. A community effort towards a knowledge-base and mathematical model of the human pathogen Salmonella Typhimurium LT2. BMC systems biology 5, 1–9 (2011).

63 Monk, J. M. et al. Genome-scale metabolic reconstructions of multiple Escherichia coli strains highlight strain-specific adaptations to nutritional environments. Proceedings of the National Academy of Sciences 110, 20338–20343 (2013).

64 Charusanti, P. et al. An experimentally-supported genome-scale metabolic network reconstruction for Yersinia pestis CO92. BMC systems biology 5, 1–13 (2011).

